# Long-term Behavioural Rewriting of Maladaptive Drinking Memories via Reconsolidation-Update Mechanisms

**DOI:** 10.1101/2020.02.06.937698

**Authors:** Grace. Gale, Vanessa E. Hennessy, Katie. Walsh, Sunjeev K. Kamboj, Ravi. K. Das

**Affiliations:** UCL; University College London

## Abstract

**Background:** Alcohol use disorders can be conceptualised as a learned pattern of maladaptive alcohol-consumption behaviours. The memories encoding these behaviours centrally contribute to long-term excessive alcohol consumption and are a key therapeutic target. The transient period of memory instability sparked during memory reconsolidation offers a therapeutic window to directly *rewrite* these memories using targeted behavioural interventions. However, clinically-relevant demonstrations of the efficacy of this approach are few. We examined key retrieval parameters for destabilising naturalistic drinking memories and the ability of subsequent counterconditioning to effect long-term reductions in drinking.

**Methods:** Hazardous/harmful beer-drinking volunteers (N=120) were factorially randomised to retrieve (RET) or not retrieve (No RET) alcohol reward memories with (PE) or without (No PE) alcohol reward prediction error. All participants subsequently underwent disgust-based *counterconditioning* of drinking cues. Acute responses to alcohol were assessed pre- and post-manipulation and drinking levels assessed up to 9 months.

**Results:** Greater long-term reductions in drinking were found when counterconditioning was conducted following retrieval (with and without PE), despite a lack of short-term group differences in motivational responding to acute alcohol. Large variability in acute levels of learning during counterconditioning were noted. ‘Responsiveness’ to counterconditioning predicted subsequent responses to acute alcohol in *RET+PE* only, consistent with reconsolidation-update mechanisms.

**Conclusions:** The longevity of behavioural interventions designed to reduce problematic drinking levels may be enhanced by leveraging reconsolidation-update mechanisms to rewrite maladaptive memory. However, inter-individual variability in levels of corrective learning is likely to determine the efficacy of reconsolidation-updating interventions and should be considered when designing and assessing interventions.

## INTRODUCTION

Harmful drinking and alcohol use disorders (AUDs) represent leading causes of global preventable mortality, contributing to 3 million deaths annually(WHO | Global status report on alcohol and health 2018 2018). This issue shows little sign of abating, with recent research suggesting an alarming increase in the prevalence of problem drinking in some demographic groups(Grant *et al.* 2017). Problematically, extant treatments for substance use disorders, including AUD, have limited long-term efficacy, with under 20% completing treatment free of dependence and fewer still maintaining abstinence for the following year (Public Health England, Department of Health, & National Drug Evidence Centre, 2018). Treatment approaches targeting the fundamental processes underlying the development and maintenance of harmful drinking are required to address this global health priority.

AUDs arise via repeated environmental exposure to alcohol amid multivariate risk factors(Sher *et al.* 2005), meaning that harmful alcohol consumption is fundamentally a *learned* pattern of maladaptive behaviours(Drummond *et al.* 1990; Hyman 2005). These behaviours arise because alcohol (like other addictive drugs) induces plasticity in mesocorticolimbic motivational circuitry(Pierce & Kumaresan 2006). This system supports reward learning, which adapts behaviour to seek out and maximise rewards when environmental cues signal that they are available. Alcohol can therefore support behavioural adaptation towards hyper-motivated alcohol seeking and consumption in the presence of environmental ‘trigger’ cues that predict alcohol availability. Practically, this manifests as arousal, and a strong desire to drink (craving) in response to certain alcohol-predictive contexts and stimuli (e.g. the sight or smell of beer) (Self 1998; Sinha & Li 2007).

To the extent that they support a harmful level of alcohol use, the memories linking environmental cues to alcohol reward can be considered to be ‘*maladaptive reward memories*’ (MRMs). Formed through repeated naturalistic exposure to alcohol and reinforced with accruing drinking episodes(Robbins *et al.* 2008), these MRMs are rapidly consolidated (McGaugh 2000), becoming highly robust and integrated into motivational networks. They display remarkable persistence (Hyman & Malenka 2001) even after extended periods of abstinence, and are therefore believed to be the core substrate underlying the persistent relapse susceptibility (even after extended abstinence) typifying AUD.

Their central pathogenic role suggests MRMs should be a primary target in the treatment of AUDs (Tronson & Taylor 2013). Current behavioural therapies may partially suppress MRMs, but do not constitute *unlearning* (Bouton 2002). As such they are a weak and temporary approach to addressing a core relapse process in AUDs. A novel approach for directly and permanently ameliorating the negative influence of MRMs on behaviour is to leverage the process of memory *reconsolidation* (Milton & Everitt 2012; Torregrossa & Taylor 2013). This is a retrieval-dependent memory maintenance process that serves to strengthen and/or update consolidated memory traces when new memory-relevant information is presented at retrieval. Such updating necessitates the temporary *destabilisation* of memory traces, such that new information can be incorporated and the relevant adjustments to the dendritic and synaptic architecture encoding the memory trace made (Clem & Huganir 2010; Merlo *et al.* 2015). If adaptive learning (for example, extinction) is timed correctly following retrieval/destabilisation, such that it occurs in the critical (∼2hour) ‘reconsolidation window’ when memories are active and unstable, it is theoretically possible to *rewrite* maladaptive memory content to a benign form (Germeroth *et al.* 2017; Monfils & Holmes 2018).

The implications of this for harmful drinking are self-evident. By directly re-formatting MRMs such that trigger cues do not provoke alcohol seeking, it may be possible to reduce alcohol consumption and prophylactically guard against relapse over the long-term. Although a nascent field, there have been highly promising early demonstrations of the potential of this approach (Walsh *et al.* 2018). Extinction (i.e. exposure therapy) following retrieval of heroin memories has been shown to produce long-lasting (6 months) reductions in craving and physiological arousal in heroin addicted patients (Xue *et al.* 2012), and reduce smoking in cigarette smokers (Germeroth *et al.* 2017). However, despite such promising early findings, there have been notable failures to replicate reconsolidation-interference effects, particularly using the retrieval-extinction paradigm (Soeter & Kindt 2011; Baker *et al.* 2013; Luyten & Beckers 2017). There are several potential interpretations for such discrepant results.

Firstly, extinction itself may represent a sub-optimal ‘corrective’ learning modality, since it is a largely passive procedure, involving no response from participants, unobserved inter-individual variability in engagement and responsiveness to extinction(Shumake *et al.* 2018) may mask effects. A promising alternative – *counterconditioning-*re-pairs cues reward cues (e.g. pictures of beer) with negatively-valenced outcomes (e.g. disgust-inducing bitter liquids and images). Disgust-counterconditioning may provide a more potent corrective learning experience than extinction (Tunstall *et al.* 2012) since it 1) leverages a potent food-rejection mechanism (Rozin & Fallon 1987) 2) the ‘disgust’ response to certain images and bitter liquids are powerful and virtually universal (Schienle *et al.* 2015) and 3) it is an ‘active’ procedure, meaning participants cannot simply disengage from the task, as occurs during extinction. We have shown broad short-term abolition of attentional biases and reactivity to alcohol cues when *counterconditioning* was conducted after MRM retrieval in hazardous drinkers(Das *et al.* 2015) a finding that has been further demonstrated in experimental animals(Goltseker *et al.* 2017), however this has never been shown to affect long-term drinking outcomes.

Secondly, memory retrieval and destabilisation are not synonymous. Retrieval procedures may fail to destabilise memories, precluding long-term rewriting. The latter explanation is compelling in the light of evidence demonstrating that memory destabilisation is highly dependent upon various ‘*boundary condition*s’(Walker & Stickgold 2016; Elsey & Kindt 2017). Primary amongst these are the *length* of retrieval (N cues presented), with retrievals that are either too short or too long failing to spark destabilisation (Suzuki *et al.* 2004; Merlo *et al.* 2014, 2018) and the presence of an appropriate ‘mismatch’ learning signal - *prediction error* (PE)(Schultz *et al.* 1997; Waelti *et al.* 2001) - at retrieval (Sevenster *et al.* 2013; Das *et al.* 2015; Krawczyk *et al.* 2017). Specifically, some level of mismatch between predicted and actual outcomes must occur to signal that the memory requires updating (Pedreira *et al.* 2004; Agustina López *et al.* 2016).

These key parameters have not been systematically manipulated in clinically-focussed reconsolidation interference studies. Most studies do not explicitly induce or test the occurrence of PE, with variable and largely heuristic retrieval lengths being employed across studies (Walsh *et al.* 2018). It is unsurprising, then, that findings are correspondingly inconsistent. In order to properly assess whether rewriting of alcohol MRMs can be reliably achieved through purely behavioural reconsolidation manipulations, a more systematic investigation of the role of MRM retrieval and prediction error prior to corrective learning is required.

In the current study, we addressed this issue by systematically manipulating MRM retrieval and the presence of prediction error at retrieval prior to a counterconditioning intervention in heavy drinkers (given the theoretical advantages of counterconditioning over extinction). We simultaneously assessed the effects of counterconditioning *per se* on cue reactivity and drinking levels and whether these were potentiated in a retrieval and prediction error-dependent manner, consistent with reconsolidation-based memory rewriting.

## METHODS

### Participants & design

120 hazardous, beer-preferring drinkers were randomised in a 2 (MRM retrieval/no retrieval) x 2 (prediction error/no prediction error) factorial design. All participants completed three sessions, corresponding to *baseline* (on Day 1), retrieval/counterconditioning *manipulation* (Day 3-5) and *post-manipulation* (Day 10 – 13). Primary inclusion criteria were : Ages 18-60, scoring >8 on the Alcohol Use Disorders Identification Test (AUDIT)(Saunders *et al.* 1993); Consuming > 40 (men) or >30 (women) UK units/week (1 unit=8g ethanol), drinking ≥4 days each week, primarily drinking beer, and having non-treatment seeking status. Exclusion criteria were: Pregnancy/breastfeeding, diagnosis of AUD/SUDs, current diagnosed psychiatric disorder, AUD as defined by the SCID; use of psychoactive medications, use of illicit drugs > 2x /month.

### Measures

#### Questionnaire assessments

The comprehensive effects of alcohol questionnaire (CEOA (Fromme *et al.* 1993)) retrospectively assessed responses to alcohol, the AUDIT, obsessive-compulsive drinking scale (OCDS (Anton *et al.* 1995)) and alcohol craving questionnaire (ACQ-NOW (Singleton *et al.* 1994)) measured maladaptive drinking patterns. Motivation to reduce drinking was measured by the stages of change readiness and treatment eagerness scale (SOCRATES (Miller & Tonigan 1996)). Distress tolerance and sensitivity to disgust were assessed by the Distress Tolerance Scale (DTS (Simons & Gaher 2005)) and Disgust Propensity and Sensitivity Scale (DPSS-R (Olatunji *et al.* 2007)), respectively. Changes in anxiety and affect due to the counterconditioning procedure were assessed using the state version of the Spielberger State-Trait Anxiety Inventory (STAI-S (Spielberger 2010)) and positive and negative affect scale (PANAS(Watson *et al.* 1988)), respectively. Drinking was quantified using the Timeline Follow-Back diary procedure(Sobell & Sobell 1992). Depressive symptomatology was assessed with the Beck Depression Inventory (BDI)(Beck *et al.* 1988).

#### Cue reactivity assessment

As in our previous study(Das *et al.* 2019), participants were presented with a 150ml glass of beer and told they would consume this after rating a series of images. They then rated their *urge to drink* and *liking of* four ‘orange juice cue’ images and four ‘beer cue’ images. These were subsequently used as retrieval cues in the ‘no retrieval’ (‘No RET’) and retrieval (‘RET’) procedures respectively on the *manipulation* day. Three *wine* and two soft drink (*neutral*) images (not used as retrieval cues) were also rated, followed by *urge to drink* the *in vivo* beer and *predicted enjoyment* of the beer. These were all rated on 11-point (0 to 10) scales. Participants then consumed the beer according to timed on-screen prompts and rated their post-consumption *actual enjoyment* of the beer and *urge to drink more* beer. These scales thus assessed the acute hedonic and motivational properties of alcohol. These *baseline* (*Day 1*) procedures both allowed assessment of changes in cue reactivity and reinforcing properties of alcohol, and set the expectation of beer consumption to maximise PE on the *manipulation* day when the drink was unexpectedly withheld in PE groups during the appropriate retrieval procedure.

#### MRM retrieval/PE procedure

was one we have previously used to reactivate alcohol MRMs and is described fully elsewhere(Das *et al.* 2015, 2019). Participants’ MRMs were retrieved by viewing/rating beer cues (*RET*). Control memories were retrieved by viewing/rating orange juice cues (*No RET*). This was identical to the cue reactivity task except 1) the *in vivo* beer was replaced with orange juice in the *No RET* groups 2) only four condition-appropriate cue images were rated. To manipulate prediction error (*PE*), the drink given to participants (orange juice or beer) was unexpectedly withheld by an on-screen prompt reading ‘*Stop, do not drink!*’ in *PE* groups: (*RET+PE* and *No RET+PE)* generating negative prediction error. In the ‘*no PE*’ conditions (*RET no PE, No RET no PE*), the drink was consumed as on *Day 1*, as expected.

#### Counterconditioning

All four groups underwent counterconditioning after the retrieval/PE manipulations as previously described(Das *et al.* 2018a). Briefly, after a 5-minute interval during which participants completed high working memory load distractor tasks (digit span, prose recall), they were shown four beer images and two neutral drink images (coffee and cola) four times each in a pseudo-randomised, fixed order. Two of the beer images (nominated ‘*Beer-Bit CSs’*) were paired with consumption of 15ml of a highly bitter solution (.067% aqueous Denatonium Benzoate/*Bitrex*). The other two beer images (nominated ‘*Beer-Pic CSs’*) were followed by one of four images taken from the IAPS database rated highly for induction of disgust. Two coffee and cola images (nominated ‘*Neut-Neut* CSs’) were followed by neutral rated images from the IAPS database. All pairings occurred on a 100% reinforcement ratio. Full information is given in the *supplementary materials*.

### Procedure

Participants responding to study advertisements were screened for eligibility by telephone. On *Day 1*, (*baseline*), participants attended UCL and completed informed consent before being breathalysed (Lion 500 Alcometer) to ensure abstinence from alcohol. They then completed demographic information (gender, age, education and smoking status) and questionnaire measures (AUDIT, Timeline follow-back, OCDS, CEOA, SOCRATES, DTS and BDI). Participants then completed the cue reactivity and acute beer rating, as described above and in the *supplementary materials*.

On *Day 2* (*manipulation: Day 1* + 48-72hrs), breath-alcohol verified abstinence was confirmed prior to completion of the DPSS-R, ACQ-NOW, PANAS and STAI. Participants then underwent group-appropriate retrieval/no-retrieval and PE/no PE manipulation followed by counterconditioning. After completion of counterconditioning participants re-completed the PANAS. On *Day 3* (*post-manipulation:* 7±2 days after *Day 2*) participants attended the test centre for the final time and recompleted all baseline questionnaires and cue reactivity/acute beer challenge before debriefing.

Remote follow-up assessments of perceived drinking changes, TLFB, ACQ-NOW and SOCRATES measures were completed at 2 weeks, 3, 6 and 9 months following *Day 3*. Participants were reimbursed at the standard university hourly rate (£10) for in-lab testing sessions and incentivised with an extra £5 for each completed remote follow-up. Sample size was calculated in G*Power 3.1.9.2 for 1-β=.95 to detect a minimum effect size of *n*_*p*_ ^*2*^*=*.05 at α=.05 for the interaction in a mixed ANOVA, assuming ρ of .5. This yielded a total required sample size of N=78 (26 per group). Anticipating minimal attrition, we randomized N=30/group.

### Statistical Approach

See *supplementary materials* for full data-handling. Changes in short-term outcomes (measured in-lab) were assessed with 2 [*Day*: *pre-manipulation* vs. *post-manipulation*) x 2 [*Retrieval: RET* vs *No RET*] x 2 [*PE*: *PE* vs No PE,] mixed ANOVA. For analysis of the cue reactivity, a factor of *Cue Type* (Beer-Bit CS/Beer-Pic CS/Neut-Neut CS/Orange Juice/Neutral) was also modelled. For counterconditioning in addition to *RET* and *PE* factors, factors of *Cue Type* (Beer-Bit CS/Beer-Pic CS/Neut-Neut CS) and *Trial* (1^st^, 2^nd^, 3^rd^, final) were included. Where sphericity was violated in repeated measures, the Greenhouse Geisser or multivariate ANOVAs were used, depending on ε values and according to published recommendations(Stevens 2012). This is reflected in multivariate/non-integer DFs.

Long-term drinking data were analysed using linear mixed models with fixed factors of *Retrieval* and *PE* across *Time* (6:Baseline, Post-manipulation, 2 weeks, 3 months, 6 months, 9 months), modelling per-participant intercepts as baseline values. *Time* slopes were initially modelled as fixed then as random, assessing improvement in model fit according to reduction >2 in Bayesian information criterion (BIC). Due to the presence of highly outlying mean daily unit alcohol consumption values for the 2-week time point (∼60 units/day, >450/week). Analyses of drinking outcomes were performed on upper-trimmed means with the trim at 30 units/day. Rating data were lost for one participant due to technical error. Alpha for all *a priori* tests was set at .05, with *p*-values Sidak-corrected for post-hoc tests. For tests of baseline trait, drinking and demographics variables, the False Discovery Rate (FDR) correction was applied. Data were analysed blind to condition.

## RESULTS

Participants were largely equivalent at baseline on key variables (see *Table 1*). Due to technical error, post-screening baseline AUDIT data were only available for *No RET no PE* N=22, *No RET+PE* N=20, *RET no PE* N=22, *RET+PE* N=20.

**Table 1:**
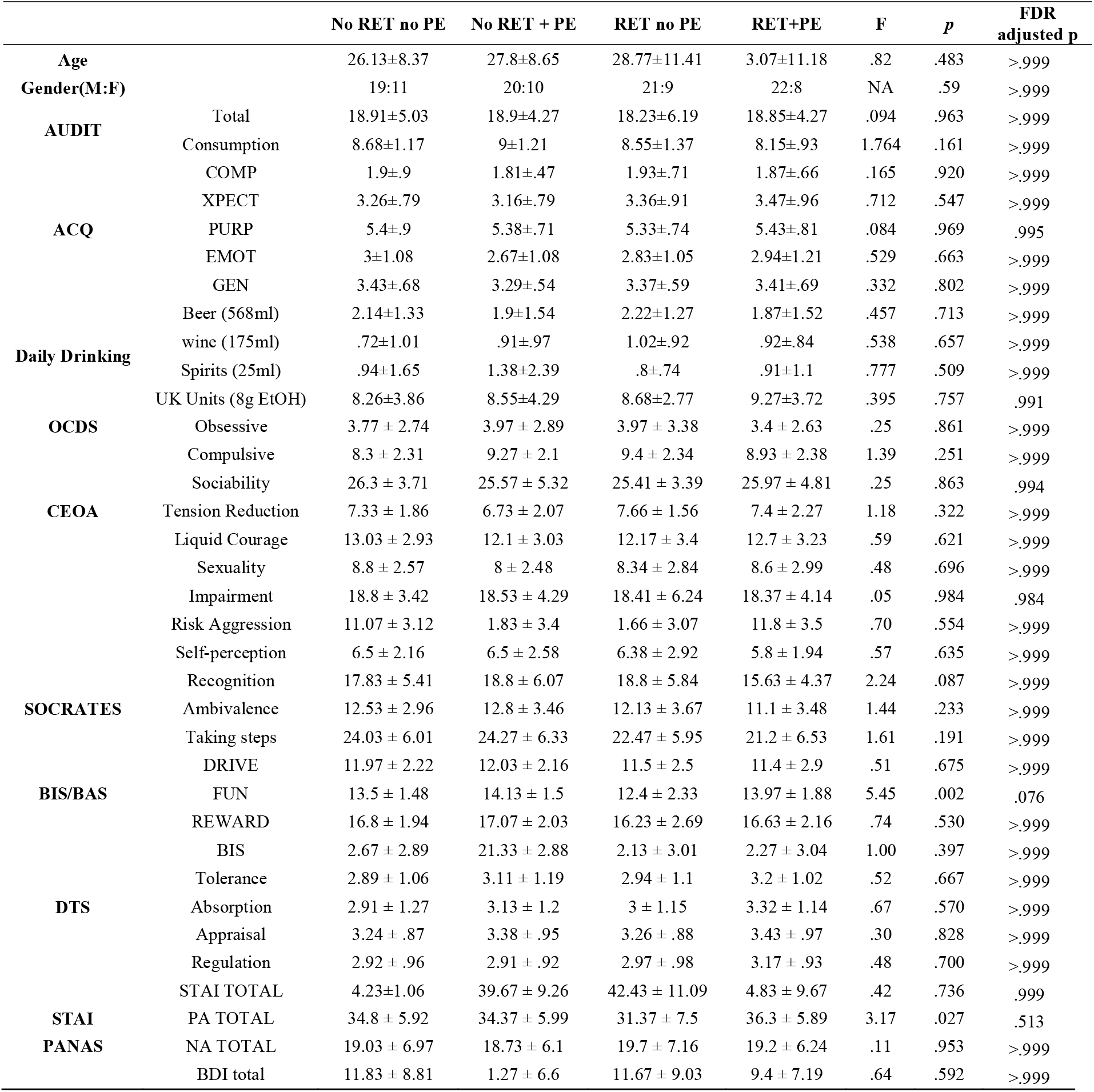
Baseline demographic drinking and questionnaire measures. Groups did not differ at false-discovery rate (FDR)-corrected alpha for any variables at baseline. Degrees of freedom for one-way ANOVA are all 3, 116, with the exception of AUDIT data where DFs were 1,83 due to data loss.

**Table 2:**
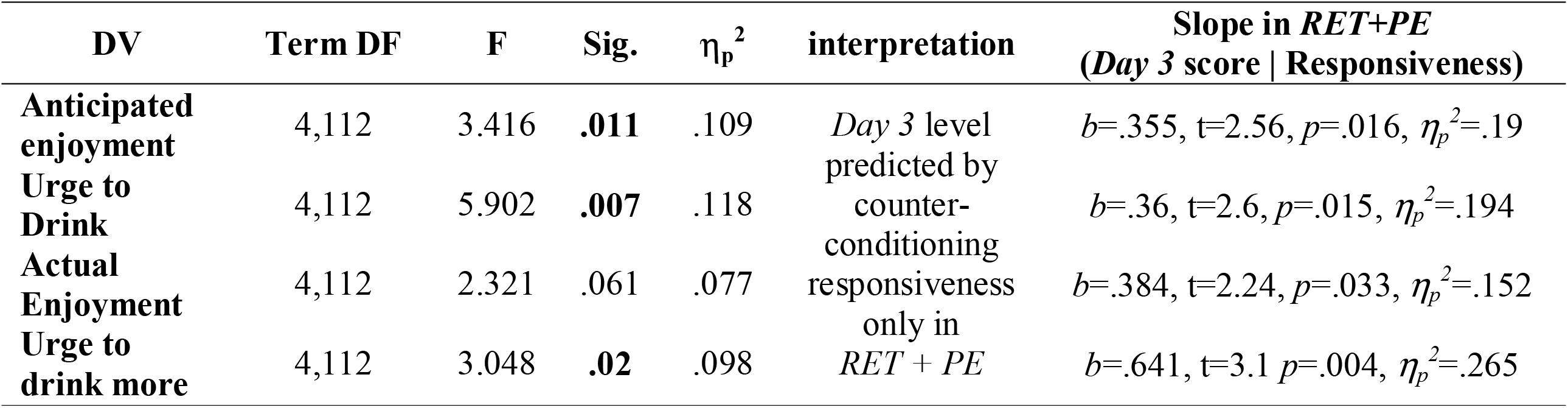
Reactivity to in-vivo beer: Highest-order (four-way) interaction terms in Day*Retrieval*Responsiveness*PE mixed ANOVAs on anticipated and actual enjoyment of sampled beer and pre and post-drink urge to drink beer. Significant effects are highlighted in bold. Degrees of freedom (DFs)=29 for all t-tests.

### Counterconditioning

A *Trial*Cue Type* interaction emerged F(4.134, 475.445)=13.656, *p*<.001, η_p_^2^=.106], indicating significant reductions in liking of Bitrex-paired beer CSs (*Beer-Bit CSs;* [F(3, 113)=19.433, *p* <.001, η_p_^2^=.34]) and disgust picture-paired beer CSs (*Beer*-*Pic CSs*; [F(3, 113)=11.274, *p*<.001, η_p_^2^=.23]) across trials, with no significant reduction in unreinforced neutral pictures (*Neut-Neut* CSs [F(3,113)=0.722, *p*=.512, *η*_*p*_^*2*^=.02]). Counterconditioning thus successfully reduced mean-level *Beer CS* liking. While successful counterconditioning was evident in both *Retrieval* groups, a marginal *Cue Type*Trial*Retrieval* interaction [F(4.134,475.445)=2.413,*η*_*p*_^*2*^=.021] indicated greater liking of *Beer-Bit CSs* [F(1,115)=6.936, *p*=.01, *η*_*p*_^*2*^=.057] and *Neut-Neut CSs* [F(1, 115)=4.594, *p=*.034, η_p_^2^=.038] in the *RET* groups vs. *No RET* groups on *Trial 1* of counterconditioning (see *Figure 1*). In the *RET* groups, all *Cue Types* were liked equally on *Trial 1* [F(1,114)=1.591, *p=*.208, η_p_^2^=.027], while in the *No RET* groups liking of *Beer-Pic CSs* was greater than *Neut-Neut CSs* [F(1,114)=9.353, *p* <.001, *η*_*p*_^*2*^=.141] on *Trial 1*. On *Trial 4* of counterconditioning, *Neut-Neut Css* were liked more than both *Beer* CSs in the *No RET* groups (*p*s≤.014) but not in the *RET* groups (*ps .072 to .956).* Unreinforced pre-exposure to CSs during MRM retrieval may have thus affected the speed and level at which these were differentiated and subsequently counterconditioned as discriminative stimuli.

**Figure 1:**
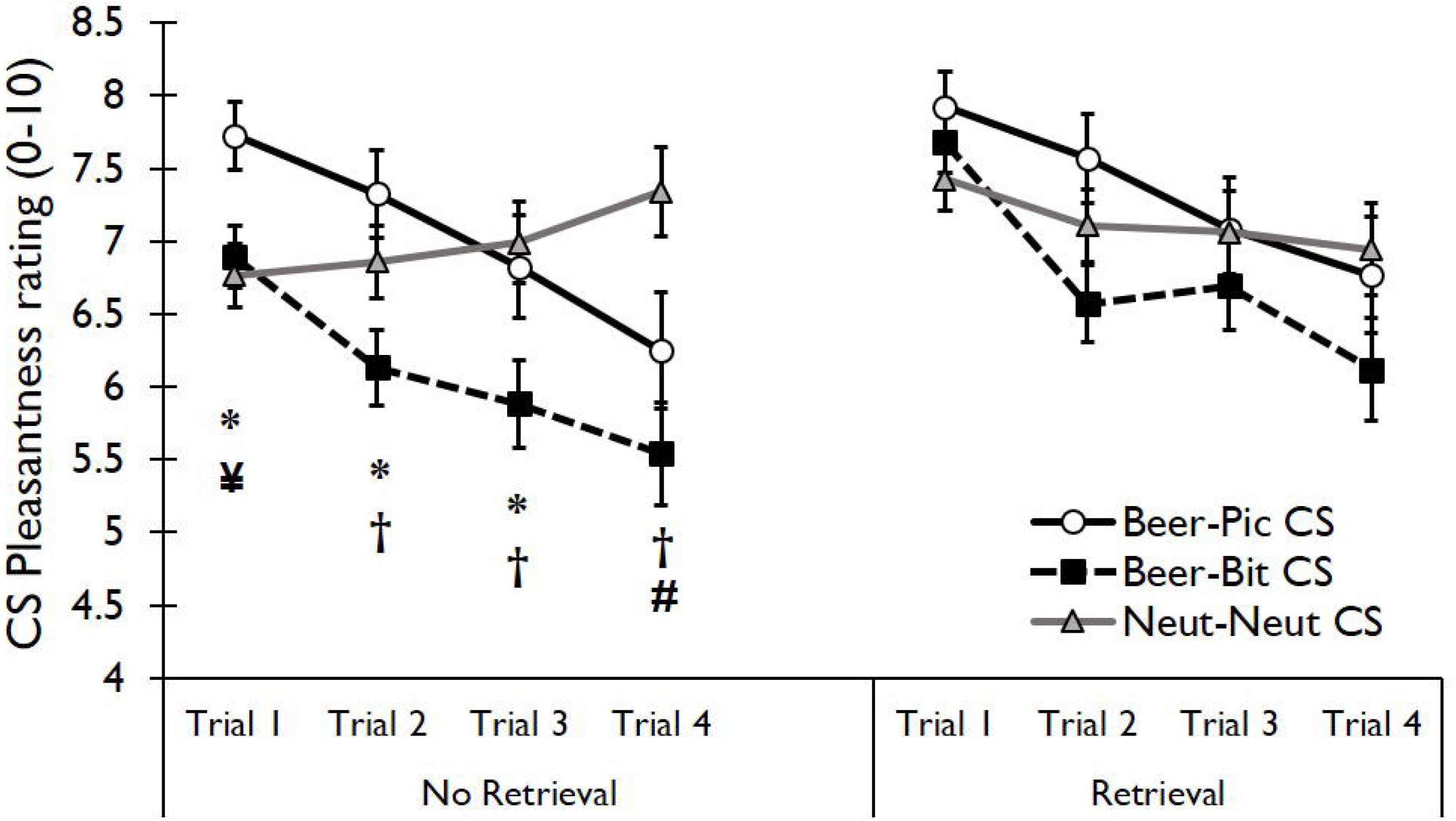
Liking ratings for the conditioned stimuli across the counterconditioning task. Significant reductions in liking of the Bitrex-paired beer CS (*Beer-Bit CS*) and disgusting image-paired beer CS (*Beer-Pic CS*) were seen in reactivated and non-reactivated groups. However only in *No RET* did the liking of CSs differ on Trial 1. **\***=*Beer-Pic*>*Neut-Neut*, **¥**=*Beer-Pic*>*Beer-Bit*, **†=***Neut-Neut*>*Beer-Bit*, **#**=*Neut-Neut*>*Beer-Pic.*

### Counterconditioning response heterogeneity

There was substantial inter-individual variation in ratings of disgust UCSs and CSs across counterconditioning. Descriptive statistics for these ratings are given in *Supplementary Table S2*. Since memory rewriting here is predicated upon level of ‘corrective learning’ (i.e. effective counterconditioning of beer cues), a measure of ‘*counterconditioning responsiveness*’ was computed as change in liking of CSs across counterconditioning (Trial 4–Trial 1). Greatest variability was seen in ratings of *Beer Pic CSs. Responsiveness* was therefore calculated as Trial 4–Trial 1 Δ in *Beer-PIC CS* liking) to be assessed as a predictor in mixed modelling of drinking outcomes and as a covariate where it was correlated with the dependent variable in general linear models (reinforcing effects of beer), including an interaction term with *Group* to assess the difference in the covariate slope across groups. Correlations with key *post-manipulation* outcomes are and exploratory analyses of trait predictors of counterconditioning responsiveness are given in *Supplementary Materials* (*Table S3*).

#### Prediction error generation

Analysis of rated ‘surprise’ levels following the retrieval and PE/no PE procedures showed a main effect of *PE*, indicating greater surprise in *PE* groups than *no PE* groups [F(1,116) = 309.79, *p*<.001, *η*_*p*_^*2*^ = .728]. This did not interact with *Retrieval* group. The PE generation procedure was thus highly successful and equally effective in *RET* and *no RET* groups. Full statistics on manipulation checks for MRM retrieval are given in the *Supplementary materials.*

### Primary Outcomes

#### Cue reactivity: Reinforcing effects of alcohol

All analyses of reinforcing effects of *in vivo* beer were analysed with *Day* (*baseline* vs. post-manipulation) x *Retrieval* (RET vs. No RET) x *PE* (PE vs. No PE) RMANCOVAs, including counterconditioning *Responsiveness* as a covariate that could interact with *RET*PE*.. Four-way interactions were found for pre-consumption *anticipated enjoyment* and *urge to drink* beer and post-consumption (primed) *urge to drink more* beer. Commensurate with the bivariate correlations, the 4-way interactions were driven *Day*Responsiveness* interactions in *RET+PE* only, indicating that degree of achieved counterconditioning predicted *post-manipulation* reactivity to *in-vivo* beer only in the ‘active’ *RET+PE* group. For *actual enjoyment* of beer (post consumption), counterconditioning responsiveness again predicted *post-manipulation* enjoyment only in *RET+* However, the 4-way interaction did not reach significance. These interaction terms and simple slopes are given in *Table 3*. Scatterplots of bivariate associations are given in *Figure 2*. Analysis of ratings of pictorial cues used in the cue reactivity task are given in the *supplementary materials*.

**Figure 2:**
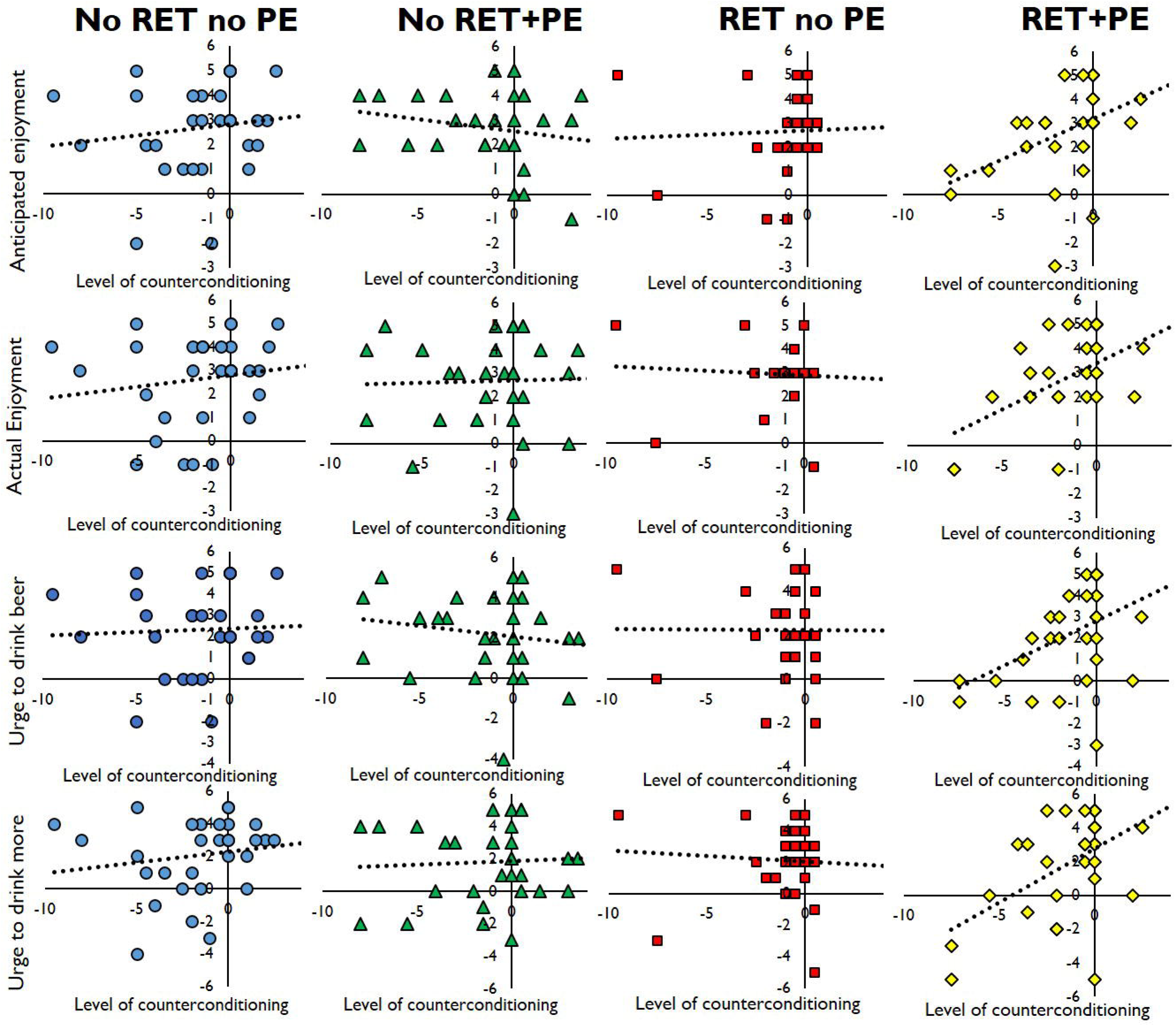
Associations between ‘strength’ of counterconditioning (change in liking of counterconditioned beer cues) anticipated enjoyment, urge to drink, actual enjoyment and urge to drink more beer on the Day 3 beer reactivity test. The correlations were significant only in RET+PE (rightmost column). Dashed lines are ordinary least-squares linear best fit lines.

### Drinking levels

#### Beer

The random intercepts-only effects mixed model revealed a significant main effect of *Time* [F(1,522.74)=39.027, *p*<.001] and a marginally significant *RET*PE*Time* interaction [F(1,522.74)=3.965, *p*=.047]. The *Time* effect represented a reduction in beer consumption across the follow-up period, with a mean reduction of .23 UK pints/day at each time point [*b*=-.232, *t*(521.5)=2.04, *p*<.0005]. The 3-way interaction represented a greater reduction in drinking across *Time* in *RET+PE* than *No RET+PE* [*b*=.146, *t*=2.06, *p*=.0397], with no differences between the other groups. Model-predicted and true values for this effect are shown below in *Figure 3* panels A and B. Modelling random slopes for *Time* did not improve model fit (BIC 2128.485→2128.919) and yielded non-significant variance in slopes (Z=1.138, *p*=.255). *Responsiveness* to counterconditioning was not a significant predictor [F(1,119.495)=.72, *p*=.679] and was detrimental to parsimonious model fit (BIC 2128.485→2134.752).

**Figure 3:**
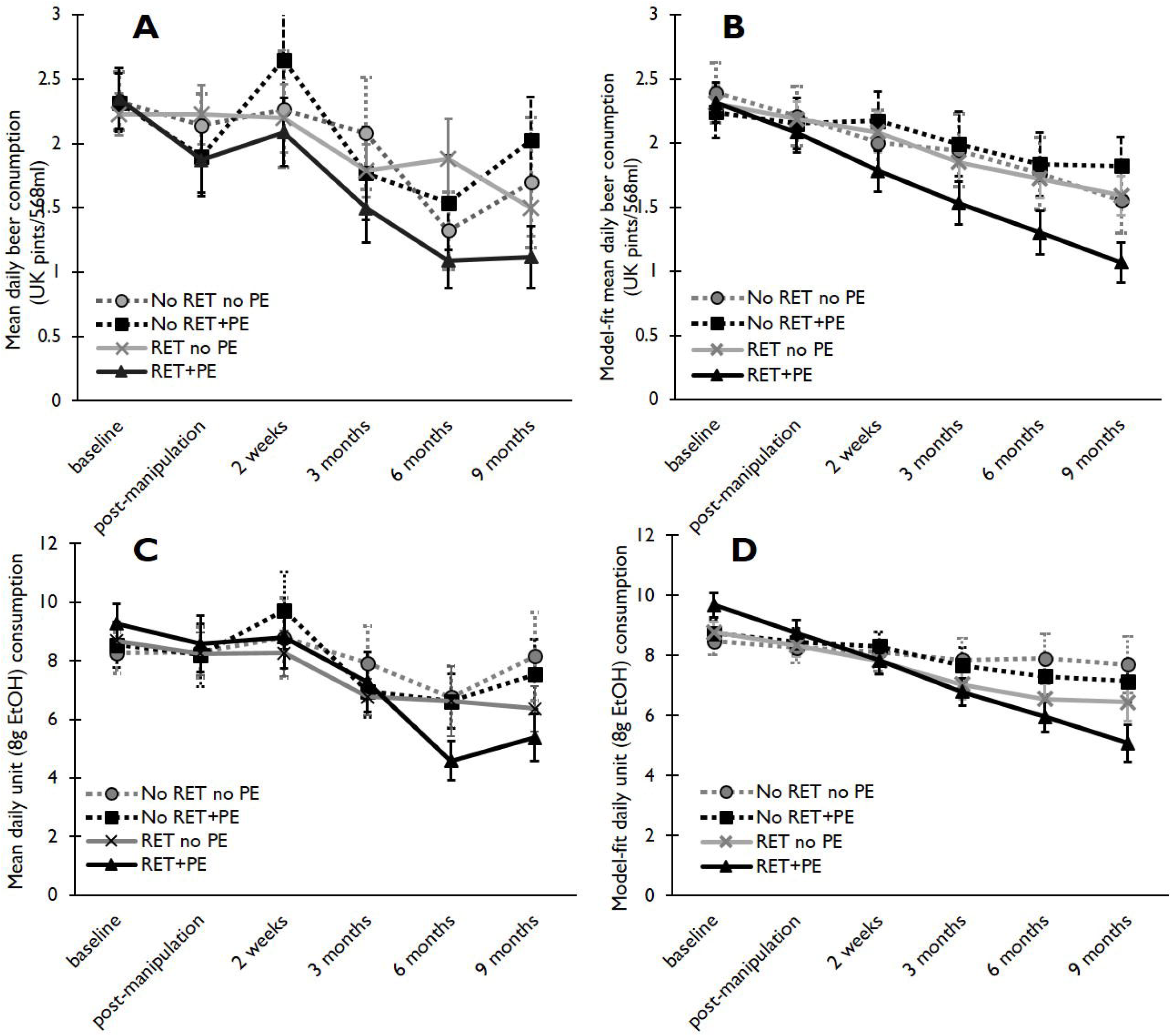
**Panel A** (top left) changes in mean daily beer consumption (in UK pints) across the study time points in each group. **Panel B (top right)** Mixed model fit values for beer consumption data. A marginally significant *Time*RET*PE* interaction indicated a steeper reduction across Time in *RET+PE* than *No RET+PE* (p=.037). **Panel C**: Changes in mean daily unit alcohol consumption across the study time points in each group. **Panel D**: Model fit values for overall alcohol consumption (total UK unit) data. A significant *RET*Time* interaction indicated significant reductions across time in RET groups but not No RET groups. Panels A&C, error bars represent SD. Panels B and D, error bars represent model SEMs.

#### Total Units

The random intercepts-only model for total unit consumption data (BIC=3748.009) yielded a significant effect of *Time* [F(1,533.775)=25.487, *p*<.001] and *RET*Time* interaction [F(1, 533.775)=4.937, *p*=.027]. Simple contrasts on the *Time* main effect against baseline drinking levels showed no overall change in drinking from baseline to post-manipulation [*b*=-.69, *t*(511.97)=.706, *p*=.48] or 2 weeks [*b=*-1.196, *t*(516.53)=.1.194, *p*=.233], with a marginal reduction by 3 months [*b=*-1.97, *t*(519.482)=1.925, *p*=.055] and significant reductions by 6 months [*b=*-4.66, *t*(519.48)=4.549, *p*<.001] and 9 months [*b*=-3.65, *t*(521.05)=3.431, *p*=.001]. Parameter estimates for the *RET*Time* interaction showed a greater reduction in drinking across *Time* in *RET* than *No RET* groups [*b=*.575, *t*(531.58)=2.192, *p*=.029]. Within-groups, the slope for the reduction in drinking across time was highly significant in the *RET* groups [*b=*-.923, *t*(51.26)=-5.008, *p*<.0005] but non-significant in the *No RET* groups [*b=*-.3, *t*(53.958)=-1.177, *p*=.245].

Significant variance in slopes [Z=2.781, *p*=.005] and improved model fit [Δ-2LL χ^2^(2)=-18.004, *p* <.001, BIC 3748.09→3743.262] when allowing slopes for *Time* to vary indicated that a random slopes effect model was appropriate. This reduced the *RET*Time* effect to only a marginally significant level [*b=*.623, *t*(107.023)=1.999, *p*=.049]. Including counterconditioning *Responsiveness* as a covariate yielded a borderline-significant predictive impact in drinking [F(1,119.518)=3.916, *p*=.05], but was detrimental to parsimonious model fit [3743.262 →3749.194], so was not included in the final model. Actual and mean model-predicted values for the *RET*Time* effect in the final model are shown in *Figure 3* panels C&D.

## DISCUSSION

We examined the potential for putative memory reconsolidation mechanisms to catalyse the efficacy and longevity of an experimental learning-based intervention in ameliorating maladaptive drinking patterns. We found mixed evidence that supported the long-term utility of a reconsolidation-focussed approach, while highlighting large response variability and potential limitations of a homogenous learning manipulation.

We observed a greater reduction in over the 9 months follow-up period when counterconditioning followed the -putatively ‘active’ *retrieval* (*RET*) *with prediction error* (PE) manipulation. Greater reductions in non-specific, *total* alcohol consumption were seen in both MRM retrieval groups, although this was not PE-dependent. These results are broadly consistent with counterconditioning updating MRMs via reconsolidation mechanisms, producing lasting beneficial changes in drinking behaviour. That lasting effects on drinking levels are observed after a one-off, purely behavioural manipulation is encouraging and extends our previous work on ketamine, suggesting reconsolidation-focussed therapies may have a bright future in the treatment of SUDs.

The current results extend our previous findings with counterconditioning during the reconsolidation window(Das *et al.* 2015) and pharmacological blockade of alcohol MRM reconsolidation by ketamine (Das et al, in press). While we previously demonstrated *RET* and *PE* –dependent beneficial effects of counterconditioning on computerised in-lab markers of MRMs, changes in responses to actual alcohol and long-term reductions in drinking following have not, until now, been shown using a purely behavioural reconsolidation-update manipulation.

Unexpectedly, the beneficial effects observed here were primarily evident only in the longer-term drinking outcomes but not acute in-lab measures of cue reactivity. The reason for this discrepancy is uncertain. One possibility is lack of sensitivity or limited ecological validity of an in-lab acute assessment of the reinforcing effects of alcohol, since anticipated enjoyment and urge to drink have no impact on whether beer is consumed or not during this test. An emergent and more compelling interpretation is that memory rewriting manipulations display their true utility when participants are exposed to naturalistic ‘high-risk’ relapse scenarios following manipulation. Indeed, previous research has also observed lagged improvements in phobic symptomatology (Soeter & Kindt 2015) and craving reductions and CO levels in smokers (Germeroth *et al.* 2017) following a reconsolidation intervention. This is in line with protection against renewal, reinstatement and spontaneous recovery conferred by reconsolidation interference in the experimental literature.

Short-term improvements are typically seen following learning-based interventions such as cue-exposure therapy, but these are not maintained across time and contexts. Indeed, in the current study, all groups largely displayed improvements in maladaptive drinking behaviours from pre–to-post-manipulation. Reconsolidation update mechanisms offer a potential means to make these interventions ‘*stick*’, vastly enhancing their long-term efficacy and protecting against relapse. The observed associations between counterconditioning responsiveness and levels of alcohol reinforcement post-manipulation in the *RET+PE* group further support this interpretation. The follow-up period used here is the longest of which we are aware in the literature. The potential for these lagged effects highlights the importance of assessing the longevity of effects over extended follow-up.

The discrepancy between retrieval *and* prediction-error-dependent effects on beer vs. all alcohol consumption was unexpected. We and others (Sevenster *et al.* 2014; Das *et al.* 2015; Exton-McGuinness *et al.* 2015; Agustina López *et al.* 2016; Krawczyk *et al.* 2017) have previously forwarded PE or ‘surprise’ at retrieval as a necessary condition for destabilisation of consolidated memories. Hypothetically, PE signals insufficient or inaccurate prediction of outcomes currently stored by the memory trace and necessitates memory destabilisation to allow the memory to update and stay ‘relevant’. These findings may seem to suggest that PE is of secondary importance in sparking memory destabilisation and reconsolidation. Indeed, most previous experimental (Milton *et al.* 2008; Saitoh *et al.* 2017; Monfils & Holmes 2018) and clinically applied (Xue *et al.* 2012, 2017; Germeroth *et al.* 2017) reconsolidation studies reporting positive findings have not explicitly manipulated prediction error. There are several key points that should be borne in mind which caution against such an interpretation, however.

It is typical in reconsolidation studies to omit the primary reinforcer during cue-driven retrieval. This will generate a variable level of PE to the extent that reinforcement is expected, despite not explicitly aiming to manipulate PE. This may well account for variability in previous findings. In the current study, although not statistically significant, the *RET+PE* group also showed the steepest overall absolute decrease in overall drinking, meaning unintended PE generation in the *RET no PE* group may have limited power to observe PE-dependent effects. Indeed, peri-retrieval ‘*surprise*’ ratings demonstrated some variability in surprise in the *RET no PE* and *RET+PE* groups, indicating that some level of unintended PE was occurring in the former group and some expectancy of deception in the latter. For clinical translation, there is minimal extra burden involved in explicitly generating and assessing PE during MRM retrieval. Indeed, in treatment scenarios (e.g. in detoxified drug-abusing patients) it would be ethically unacceptable to reinforce patients with abused drugs. Moreover, there are no demonstrations of *inferiority* of PE vs. no PE at retrieval in memory destabilisation. The most prudent course of action would be to include PE-generation procedures in experimental and translational retrieval protocols going forward and at the very least assess these explicitly.

### Limitations

We have previously assumed a relatively homogenous response to the counterconditioning intervention, given that is leverages very basic learning and aversion mechanisms. The large observed variability in level of achieved counterconditioning or ‘responsiveness’ demonstrate that this assumption is not tenable. Some participants displayed reductions of in liking of negatively reinforced beer stimuli over half the scale range while others showed little or no change and some even displayed *increased* liking over the course of the task. Equally, some participants did not rate the UCSs as particularly aversive, with some even rating them as mildly pleasant. Having extensively piloted the doses of Bitrex used here ourselves, this is puzzling to us, although genetic polymorphisms moderating bitterness perception may play a key role(Duffy & Bartoshuk 2000). We further found that disgust propensity, sensitivity and distress tolerance predicted counterconditioning responsiveness, yielding potentially useful trait markers of likely treatment response. However, such individual variability to counterconditioning likely obscured potential group-level differences in responses to the acute alcohol challenge.

One could reasonably anticipate equal (or greater) response variability when using retrieval-extinction (Shumake *et al.* 2018); a paradigm that has dominated behavioural memory rewriting research. This may partially explain the inconsistencies and difficulties replicating findings with retrieval-extinction interventions(Soeter & Kindt 2011; Baker *et al.* 2013; Chen *et al.* 2014; Luyten & Beckers 2017), since a failure to extinguish would preclude any potentiating effect of prior memory retrieval. These observations highlight the importance assessing level of corrective learning, conducting learning to a criterion level or identifying potential low-responders within reconsolidation-updating paradigms.

Variability in learning is perhaps a reason to recommend pharmacological memory-weakening over purely behavioural memory updating approaches in certain populations. Drugs’ pharmacodynamic profiles are generally not subject to influence by individual cognitive variables like learning rates, boredom and punishment insensitivity and may be a key option where behavioural approaches fail.

There is no way of assessing whether the *RET+PE* truly destabilised alcohol MRMs and engaged reconsolidation mechanisms (or did so to an equal degree) in all individuals in the current study, since memory destabilisation is a behaviourally silent process. This remains the primary impediment to translational/clinical developments within the reconsolidation field, which is in desperate need of validated biomarkers of memory destabilisation. The lack of triangulation between short-term lab measures and longer-term drinking outcomes compounds this issue in the current study. We have, however, now demonstrated group-level sufficiency of the *RET+PE* procedure used improving clinically-relevant outcomes in five studies (Das *et al.* 2015, 2018a, 2018b, 2019; Hon *et al.* 2016). Along with the apparently durable effects on drinking observed here, this lends support to the notion that reconsolidation mechanisms were engaged in the current study. While non reconsolidation mechanisms may explain shorter-term effects on outcome, the emergence of divergent effects longer-term observed here are in line with reconsolidation-update.

The current study highlights fundamental questions regarding the parameters that conspire retrieval conspire to determine the fate of memories at retrieval. The future of memory-rewriting interventions will rely upon better understanding of these parameters and individual optimisation memory destabilisation procedures based therein. Nevertheless, the results obtained here are should energise future research in the field.

## Financial Support

This work was funded by a Medical Research Council grant awarded to Sunjeev Kamboj (grant number: MR/M007006/1)

## Conflicts of interest

None

## Ethical Standards

*The authors assert that all procedures contributing to this work comply with the ethical standards of the relevant national and institutional committees on human experimentation and with the Helsinki Declaration of 1975, as revised in 2013.*

---

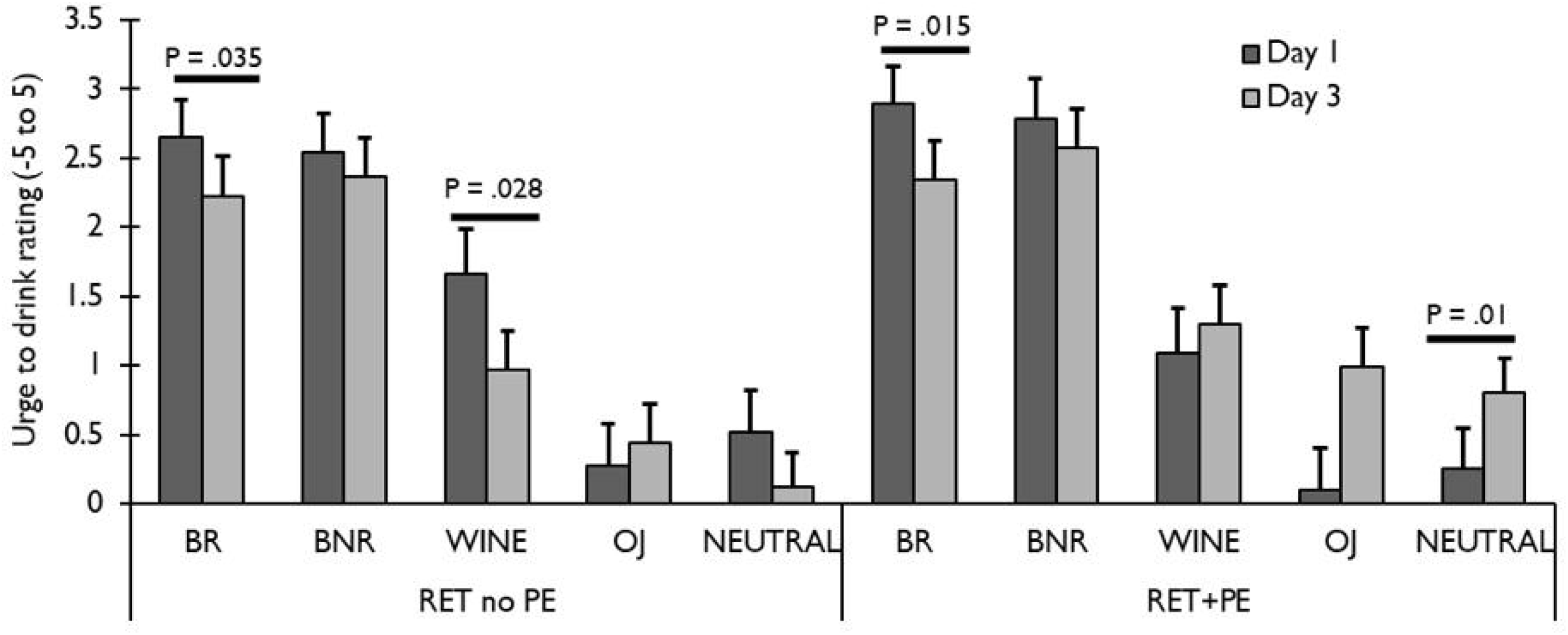

## REFERENCES

Agustina López M, Jimena Santos M, Cortasa S, Fernández RS, Carbó Tano M, Pedreira ME (2016). Different dimensions of the prediction error as a decisive factor for the triggering of the reconsolidation process. Neurobiology of Learning and Memory 136, 210–219.

Anton RF, Moak DH, Latham P (1995). The Obsessive Compulsive Drinking Scale: A Self-Rated Instrument for the Quantification of Thoughts about Alcohol and Drinking Behavior. Alcoholism: Clinical and Experimental Research 19, 92–99.

Baker KD, McNally GP, Richardson R (2013). Memory retrieval before or after extinction reduces recovery of fear in adolescent rats. Learning and Memory

Beck AT, Steer RA, Carbin MG (1988). Psychometric properties of the Beck Depression Inventory: Twenty-five years of evaluation. Clinical psychology review 8, 77–100.

Bouton ME (2002). Context, ambiguity, and unlearning: sources of relapse after behavioral extinction. Biological psychiatry 52, 976–986.

Chen S, Cai D, Pearce K, Sun PYW, Roberts AC, Glanzman DL (2014). Reinstatement of long-term memory following erasure of its behavioral and synaptic expression in Aplysia. eLife 3, e03896.

Clem RL, Huganir RL (2010). Calcium-permeable AMPA receptor dynamics mediate fear memory erasure. Science 330, 1108–1112.

Das RK, Gale G, Hennessy V, Kamboj SK (2018a). A Prediction Error-driven Retrieval Procedure for Destabilizing and Rewriting Maladaptive Reward Memories in Hazardous Drinkers. Journal of Visualized Experiments 56097, e56097–e56097.

Das RK, Gale G, Walsh K, Hennessy VE, Iskandar G, Mordecai LA, Brandner B, Kindt M, Curran HV, Kamboj SK (2019). Ketamine can reduce harmful drinking by pharmacologically rewriting drinking memories. Nature Communications 10, 5187.

Das RK, Lawn W, Kamboj SK (2015). Rewriting the valuation and salience of alcohol-related stimuli via memory reconsolidation.. Nature Publishing Group Translational Psychiatry 5, e645–e645.

Das RK, Walsh K, Hannaford J, Lazzarino AI, Kamboj SK (2018b). Nitrous oxide may interfere with the reconsolidation of drinking memories in hazardous drinkers in a prediction-error-dependent manner.. Elsevier European Neuropsychopharmacology 28, 828–840.

Drummond DC, Cooper T, Glautier SP (1990). Conditioned learning in alcohol dependence: implications for cue exposure treatment. British journal of addiction 85, 725–743.

Duffy VB, Bartoshuk LM (2000). Food acceptance and genetic variation in taste.. Elsevier Journal of the American Dietetic Association 100, 647–655.

Elsey JWB, Kindt M (2017). Breaking boundaries: Optimizing reconsolidation-based interventions for strong and old memories. Learning & Memory (Cold Spring Harbor, N.Y.) 24, 472–479.

Exton-McGuinness MTJ, Lee JLC, Reichelt AC (2015). Updating memories-The role of prediction errors in memory reconsolidation. Behavioural Brain Research 278

Fromme K, Stroot EA, Kaplan D (1993). Comprehensive effects of alcohol: Development and psychometric assessment of a new expectancy questionnaire. Psychological Assessment 5, 19–26.

Germeroth LJ, Carpenter MJ, Baker NL, Froeliger B, LaRowe SD, Saladin ME (2017). Effect of a Brief Memory Updating Intervention on Smoking Behavior. JAMA Psychiatry 74, 214.

Goltseker K, Bolotin L, Barak S (2017). Counterconditioning During Reconsolidation Prevents Relapse of Cocaine Memories. American College of Neuropsychopharmacology Neuropsychopharmacology 42, 716–726.

Grant BF, Chou SP, Saha TD, Pickering RP, Kerridge BT, Ruan WJ, Huang B, Jung J, Zhang H, Fan A, Hasin DS (2017). Prevalence of 12-Month Alcohol Use, High-Risk Drinking, and DSM-IV Alcohol Use Disorder in the United States, 2001-2002 to 2012-2013. JAMA Psychiatry 74, 911.

Hon T, Das RKRK, Kamboj SKSKSK (2016). The effects of cognitive reappraisal following retrieval-procedures designed to destabilize alcohol memories in high-risk drinkers.. Springer Berlin Heidelberg Psychopharmacology 233, 851–861.

Hyman SE (2005). Addiction: a disease of learning and memory. American Journal of Psychiatry 162, 1414–1422.

Hyman SE, Malenka RC (2001). Addiction and the brain: the neurobiology of compulsion and its persistence. Nature Reviews Neuroscience 2, 695–703.

Krawczyk MC, Fernández RS, Pedreira ME, Boccia MM (2017). Toward a better understanding on the role of prediction error on memory processes: From bench to clinic.. Academic Press Neurobiology of Learning and Memory 142, 13–20.

Luyten L, Beckers T (2017). A preregistered, direct replication attempt of the retrieval-extinction effect in cued fear conditioning in rats. Neurobiology of Learning and Memory

McGaugh JL (2000). Memory--a Century of Consolidation. Science 287, 248–251.

Merlo E, Bekinschtein P, Jonkman S, Medina JH (2015). Molecular Mechanisms of Memory Consolidation, Reconsolidation, and Persistence. Neural Plasticity 2015, 1–2.

Merlo E, Milton AL, Everitt BJ (2018). A Novel Retrieval-Dependent Memory Process Revealed by the Arrest of ERK1/2 Activation in the Basolateral Amygdala.. Society for Neuroscience The Journal of Neuroscience 38, 3199–3207.

Merlo E, Milton AL, Goozée ZY, Theobald DE, Everitt BJ (2014). Reconsolidation and Extinction Are Dissociable and Mutually Exclusive Processes: Behavioral and Molecular Evidence. The Journal of Neuroscience 34, 2422–2431.

Miller WR, Tonigan JS (1996). Assessing drinkers’ motivation for change: The Stages of Change Readiness and Treatment Eagerness Scale (SOCRATES). Psychology of Addictive Behaviors 10, 81–89.

Milton AL, Everitt BJ (2012). The persistence of maladaptive memory: addiction, drug memories and anti-relapse treatments.. Elsevier Ltd Neuroscience & Biobehavioral Reviews 36, 1119–1139.

Milton AL, Lee JLC, Butler VJ, Gardner R, Everitt BJ (2008). Intra-amygdala and systemic antagonism of NMDA receptors prevents the reconsolidation of drug-associated memory and impairs subsequently both novel and previously acquired drug-seeking behaviors. The Journal of Neuroscience 28, 8230–8237.

Monfils MH, Holmes EA (2018). Memory boundaries: Opening a window inspired by reconsolidation to treat anxiety, trauma-related, and addiction disorders.. Elsevier Science The Lancet Psychiatry 5, 1032–1042.

Olatunji BO, Cisler JM, Deacon BJ, Connolly K, Lohr JM (2007). The Disgust Propensity and Sensitivity Scale-Revised: Psychometric properties and specificity in relation to anxiety disorder symptoms. Journal of Anxiety Disorders 21, 918–930.

Pedreira ME, Pérez-Cuesta LM, Maldonado H (2004). Mismatch between what is expected and what actually occurs triggers memory reconsolidation or extinction. Learning & Memory 11, 579–585.

Pierce RC, Kumaresan V (2006). The mesolimbic dopamine system: The final common pathway for the reinforcing effect of drugs of abuse? Neuroscience & Biobehavioral Reviews 30, 215–238.

Public HealthEngland, Department of Health, National Drug Evidence Centre (2018). Adult Drug Statistics from the National Drug Treatment Monitoring System (NDTMS), 38.

Robbins TW, Ersche KD, Everitt BJ (2008). Drug addiction and the memory systems of the brain. Annals of the New York Academy of Sciences 1141, 1–21.

Rozin P, Fallon AE (1987). A Perspective on Disgust. Psychological Review

Saitoh A, Akagi K, Oka J-I, Yamada M (2017). Post-reexposure administration of d-cycloserine facilitates reconsolidation of contextual conditioned fear memory in rats.. Springer Journal of Neural Transmission 124, 583–587.

Saunders JB, Aasland OG, Babor TF, Delafuente JR, Grant M, De La Fuente JR, Grant M, Delafuente JR, Grant M (1993). Development of the Alcohol Use Disorders Identification Test (AUDIT): WHO Collaborative Project on Early Detection of Persons with Harmful Alcohol Consumption-II. Addiction 88, 791–804.

Schienle A, Arendasy M, Schwab D (2015). Disgust Responses to Bitter Compounds: the Role of Disgust Sensitivity. Chemosensory Perception 8, 167–173.

Schultz W, Dayan P, Montague PR (1997). A neural substrate of prediction and reward. Science 275, 1593–1599.

Self DW (1998). Neural substrates of drug craving and relapse in drug addiction. Annals of Medicine 30, 379–389.

Sevenster D, Beckers T, Kindt M (2013). Prediction error governs pharmacologically induced amnesia for learned fear. Science 339, 830–833.

Sevenster D, Beckers T, Kindt M (2014). Prediction error demarcates the transition from retrieval, to reconsolidation, to new learning. Learning & Memory 21, 580–584.

Sher KJ, Grekin ER, Williams NA (2005). The Development of Alcohol Use Disorders.. Annual Reviews Annual Review of Clinical Psychology 1, 493–523.

Shumake J, Jones C, Auchter A, Monfils M-H (2018). Data-driven criteria to assess fear remission and phenotypic variability of extinction in rats.. The Royal Society Philosophical Transactions of the Royal Society B: Biological Sciences 373, 20170035.

Simons JS, Gaher RM (2005). The Distress Tolerance Scale: Development and Validation of a Self-Report Measure. Motivation and Emotion 29, 83–102.

Singleton EG, Henningfield JE, Tiffany ST (1994). Alcohol craving questionnaire: ACQ-Now: background and administration manual. Baltimore: NIDA Addiction Research Centre

Sinha R, Li CSR (2007). Imaging stress- and cue-induced drug and alcohol craving: association with relapse and clinical implications. Drug and Alcohol Review 26, 25–31.

Sobell LC, Sobell MB (1992). Timeline follow-back. In Measuring alcohol consumption, pp 41–72. Springer.

Soeter M, Kindt M (2011). Disrupting reconsolidation: Pharmacological and behavioral manipulations. Learning & Memory 18, 357–366.

Soeter M, Kindt M (2015). An abrupt transformation of phobic behavior after a post-retrieval amnesic agent.. Elsevier Biological psychiatry 78, 880–886.

Spielberger CD (2010). State□Trait Anxiety Inventory. Wiley Online Library.

Stevens JP (2012). Applied multivariate statistics for the social sciences. Routledge.

Suzuki A, Josselyn SA, Frankland PW, Masushige S, Silva AJ, Kida S (2004). Memory reconsolidation and extinction have distinct temporal and biochemical signatures. The Journal of Neuroscience 24, 4787–4795.

Torregrossa MM, Taylor JR (2013). Learning to forget: manipulating extinction and reconsolidation processes to treat addiction. Psychopharmacology 226, 659–672.

Tronson NC, Taylor JR (2013). *Addiction: A drug-induced disorder of memory reconsolidation*.. Elsevier Current Trends Current Opinion in Neurobiology 23, 573–580.

Tunstall BJ, Verendeev A, Kearns DN (2012). A comparison of therapies for the treatment of drug cues: Counterconditioning vs. extinction in male rats. Experimental and Clinical Psychopharmacology 20, 447–453.

Waelti P, Dickinson A, Schultz W (2001). Dopamine responses comply with basic assumptions of formal learning theory. Nature 412, 43–48.

Walker MP, Stickgold R (2016). Understanding the boundary conditions of memory reconsolidation. Proceedings of the National Academy of Sciences of the United States of America 113, E3991–E3992.

Walsh KH, Das RK, Saladin ME, Kamboj SK (2018). Modulation of naturalistic maladaptive memories using behavioural and pharmacological reconsolidation-interfering strategies: a systematic review and meta-analysis of clinical and ‘sub-clinical’ studies.. Springer Berlin Heidelberg Psychopharmacology 235, 2507–2527.

Watson D, Clark LA, Tellegen A (1988). Development and validation of brief measures of positive and negative affect: The PANAS scales. Journal of Personality and Social Psychology 54, 1063–1070.

WHO | Global status report on alcohol and health 2018 (2018).. World Health Organization WHO

Xue Y-X, Deng J-H, Chen Y-Y, Zhang L-B, Wu P, Huang G-D, Luo Y-X, Bao Y-P, Wang Y-M, Shaham Y, Shi J, Lu L (2017). Effect of selective inhibition of reactivated nicotine-associated memories with propranolol on nicotine craving.. American Medical Association JAMA Psychiatry 74, 224–232.

Xue Y-XY-X, Luo Y-X, Wu P, Shi H-SH-S, Xue L-F, Chen C, Zhu W-LW-L, Ding Z-BZ-B, Bao YY -p., Shi J, Epstein DH, Shaham Y, Lu L (2012). A Memory Retrieval-Extinction Procedure to Prevent Drug Craving and Relapse. Science 336, 241–245.

