## Supplementary material for "Long-term Behavioural Rewriting of Maladaptive Drinking Memories via Reconsolidation-Update Mechanisms": Supplemetal Table 1

*Table S1*: N respondents in each group at each time point from baseline to final follow up for all drinking-related measures.

|  |  | baseline | post-manipulation | 2 weeks | 3 months | 6 months | 9 months |
| --- | --- | --- | --- | --- | --- | --- | --- |
| AUDIT | No RET no PE | 22 | 30 | 27 | 25 | 25 | 26 |
|  | No RET+PE | 20 | 29 | 24 | 23 | 23 | 26 |
|  | RET no PE | 22 | 30 | 27 | 23 | 23 | 23 |
|  | RET+PE | 20 | 29 | 30 | 27 | 27 | 23 |
| TLFB | No RET no PE | 30 | 30 | 27 | 23 | 23 | 26 |
|  | No RET+PE | 30 | 30 | 24 | 21 | 22 | 25 |
|  | RET no PE | 30 | 30 | 27 | 23 | 23 | 23 |
|  | RET+PE | 30 | 30 | 29 | 26 | 26 | 23 |
| SOCRATES | No RET no PE | 30 | 30 | 27 | 25 | 26 | 26 |
|  | No RET+PE | 30 | 30 | 24 | 23 | 24 | 26 |
|  | RET no PE | 30 | 30 | 27 | 23 | 22 | 23 |
|  | RET+PE | 30 | 30 | 30 | 27 | 25 | 22 |
| ACQ | No RET no PE | 30 | 30 | 27 | 25 | 26 | 26 |
|  | No RET+PE | 30 | 30 | 24 | 23 | 24 | 26 |
|  | RET no PE | 30 | 30 | 27 | 23 | 22 | 23 |
|  | RET+PE | 30 | 30 | 30 | 27 | 25 | 22 |
