## Supplementary material for "Long-term Behavioural Rewriting of Maladaptive Drinking Memories via Reconsolidation-Update Mechanisms": Supplemetal Table 1

*Table S2:* Variability in responses to CSs and UCSs during counterconditioning. Response heterogeneity in ‘level’ of counterconditioning is evident in the range of liking ratings and standard deviation (SD).

|  | **Min** | **Max** | **Mean** | **SD** |
| --- | --- | --- | --- | --- |
| *Beer-Pic CS liking Trial 1* | 2.5 | 10 | 7.82 | 1.82 |
| *Beer-Pic CS liking Last Trial* | 0 | 10 | 6.51 | 3.07 |
| *Beer-Bit CS liking Trial 1* | 2.5 | 10 | 7.28 | 1.68 |
| *Beer-Bit CS liking Last Trial* | 0 | 10 | 5.83 | 2.71 |
| *Neut-Neut CS liking Trial 1* | 2.5 | 10 | 7.1 | 1.71 |
| *Neut-Neut CS liking last Trial* | 0 | 10 | 7.14 | 2.39 |
| *∆ Beer-Pic CS liking* | -9.5 | 4.5 | -1.3 | 2.67 |
| *∆ Beer-Bit CS liking* | -9 | 3 | -1.44 | 2.47 |
| *∆ Neut-Neut CS liking* | -9.5 | 5.5 | 0.05 | 2.25 |
| *Bitrex UCS liking Trial 1* | 0 | 6 | 1.58 | 1.57 |
| *Bitrex UCS liking Last Trial* | 0 | 6.5 | 1.25 | 1.63 |
| *Pic UCS liking Trial 1* | 0 | 10 | 1.58 | 1.85 |
| *Pic UCS liking Last Trial* | 0 | 10 | 1.61 | 1.89 |
| *Neut UCS liking Trial 1* | 0 | 10 | 5.23 | 2.04 |
| *Neut UCS liking Last Trial* | 0 | 10 | 5.76 | 2.15 |
