## Supplementary material for "Long-term Behavioural Rewriting of Maladaptive Drinking Memories via Reconsolidation-Update Mechanisms": Supplemetal Table 3

*Table S3*: Pearson’s correlations between acute changes in liking of beer cues counter conditioned with Bitrex (*Beer-Bit*) and pictorial (Beer-Pic) UCSs with Day 3 cue and alcohol reactivity outcomes.

|  |  | **No RET No PE** | | **No RET + PE** | | **RET no PE** | | **RET+PE** | |
| --- | --- | --- | --- | --- | --- | --- | --- | --- | --- |
|  |  | ∆ Beer-BIT | ∆ Beer-PIC | ∆ Beer-BIT | ∆ Beer-PIC | ∆ Beer-BIT | ∆ Beer-PIC | ∆ Beer-BIT | ∆ Beer-PIC |
| **Cue image Ratings** | **Beer-React liking** | .089 | -.101 | -.188 | -.106 | -.113 | .21 | .178 | .212 |
|  | **Beer-Non-React liking** | -.086 | -.121 | -.1 | -.025 | -.084 | .161 | .185 | **.37*** |
|  | **Wine Liking** | .085 | .141 | -.023 | -.02 | -.075 | .278 | .275 | .322 |
|  | **OJ Liking** | .249 | -.131 | -.149 | -.20 | -.096 | -.117 | -.204 | -.071 |
|  | **Beer-React urge** | .043 | .073 | -.242 | -.161 | .249 | -.013 | .171 | .295 |
|  | **Beer-Non-React urge** | -.087 | -.142 | -.154 | -.099 | .192 | -.012 | .222 | .319 |
|  | **Wine Urge** | .08 | .184 | -.093 | -.026 | -.101 | -.315 | .134 | **.39*** |
|  | **OJ Urge** | .093 | -.155 | -.177 | -.083 | .225 | -.177 | .077 | **.38*** |
| **In vivo beer ratings** | **Drink itself liking** | -.016 | .066 | -.21 | -.173 | -.154 | .101 | .324 | .058 |
|  | **Drink itself urge** | .037 | .086 | -.314 | -.221 | -.262 | .155 | **.363*** | **.441*** |
|  | **anticipated enjoyment** | .137 | .074 | -.243 | -.188 | -.35 | .052 | .247 | **.436*** |
|  | **Anticipatory urge** | .046 | .046 | -.314 | -.159 | -.349 | -.006 | .31 | **.445*** |
|  | **Drink enjoyment** | .148 | .209 | -.026 | .027 | -.143 | -.073 | .305 | **.39*** |
|  | **Post-drink want more** | .161 | .098 | -.041 | .053 | -.016 | -.075 | .**367*** | **.515*** |
