## Supplementary file for "Long-term Behavioural Rewriting of Maladaptive Drinking Memories via Reconsolidation-Update Mechanisms"

**SUPPLEMENTARY MATERIALS**

**METHODS**

**Counterconditioning trial information:**

On each trial, a ‘cue image’ (CS) was presented alone for 10 second on the left side of the screen in a 400x400 pixel square. This was followed the ‘outcome’ unconditioned stimulus (UCS). Two negatively reinforcing UCSs were used. The first was 15ml of a 0.067% aqueous solution of denatonium benzoate (Bitrex). This is an extremely bitter solution that reliably produces disgust responses. The second UCS type consisted of four images rated highly for disgust, sourced from the IAPS database. Two of the beer images used as CSs were designated ‘*Beer Bit CSs*’ and would be paired with the Bitrex UCS four times each. The remaining two beer images were designated *Beer Pic CSs* and paired once each with each of the four disgust-induction images from the IAPS database. The designation of beer images to as *Beer Pic* or *Beer*-*Bit CSs* was random. To control for non-associative effects, two soft drink images were designated ‘*neutral*’ cues and paired with affectively neutral images of office furniture taken from the IAPS database. As both CSs and outcomes in these trials were neutral, they were designated ‘*Neut-Neut CSs’*.

On *Beer-Bitrex CS* trials, this was a screen saying ‘*Drink Now’*, prompting consumption of the Bitrex UCS. Eight Bitrex (Bit) UCSs were delivered in total in opaque paper cups. Participants were required to drink all of the liquid in the cup before moving on to the next trial. The remaining number of cups of the Bitrex UCS was unknown by the participant, with the cups themselves stored behind a screen. On *Beer-Pic CS* and *Neut-Neut CS* trials, this was the disgusting or neutral UCS image displayed for 10 seconds, as appropriate. On each trial, the CS image appeared for ten seconds during which time participants participants rated the CS’s pleasantness. The ‘outcome’ UCS then appeared for another ten seconds while participants either looked at the outcome image (*Beer-Pic* and *Neut Neut* *CS trials*) or drank the Bitrex solution. All images then disappeared and a rating scale for the UCS’s pleasantness appeared. All pleasantness ratings were on a scale from 0 (extremely unpleasant) to 10 (extremely pleasant). Counterconditioning was 24 trials in total, consisting of 8 *Beer-Bitrex C*S trials, 8 *Beer*-*Pic CS* trials and 8 *Neut-Neut CS* trials. Trial types were presented in a pseudo-randomised order with the constraints that no more than two of each type of CS could appear for more than two trials consecutively. Following counterconditioning, all participants were given a square of milk chocolate to mitigate the taste of Bitrex.

**Statistical Approach and data handling**

***Statistical Approach:***

Data analysis was performed using IBM SPSS 25 for Windows. Where sphericity was violated in repeated measures, the Greenhouse Geisser correction or multivariate terms were used, depending on ε values and according to published recommendations^60^. This is reflected in non-integer DFs in reported ANOVAs. Changes in short-term drinking-related dependent variables (measure in-lab) were assessed with 2 x 2 x 2mixed ANOVA: within-subjects factor = *Day* (pre-manipulation vs. post-manipulation), between-subjects factors = *Retrieval* (RET vs No RET) and PE (PE, no PE). For analysis of the counterconditioning task, factors of *Cue Type* (Beer-Bit CS/ Beer-Pic CS/ Neut-Neut CS) and *Trial* (1^st^, 2^nd^, 3^rd^, final) were included. The four levels of the *Trial* factor were calculated by taking the mean ratings of each two consecutive presentation of each *CS Type*. Significant interactions in omnibus were investigated with multivariate simple effects analyses and paired tests on marginal means, where appropriate.

Long-term drinking levels were (mean daily beer consumption, mean daily UK units) were analysed using linear mixed models with fixed factors of *Retrieval* and *PE* across *Time* (6:Baseline, Post-manipulation, 2 weeks, 3 months, 6 months, 9 months), modelling per-participant random intercepts as baseline values. *Time* slopes were initially modelled as fixed, with all factorial interactions then allowed to vary randomly, assessing improvement in model fit according to reduction in Bayesian information criterion (BIC) and chi-square tests on -2 log likelihood (-2LL). A reduction >2 in BIC represents an improvement in complexity-penalised model fit. Mixed models were estimated using maximum likelihood with unstructured working correlation matrices. Due to the presence of a small number of unfeasibly high, outlying mean weekly beer consumption values (> 60 units per day, > 400 units/week), analyses were performed on upper-trimmed means with the trim point set at/above 30 units/day. This successfully removed the outlying values. Rating data during counterconditioning were lost for one participant due to technical error. Alpha for all *a priori* tests was set at 0.05, with *p*-values Bonferroni- corrected for post-hoc tests. For tests of baseline trait, drinking and demographics difference, the False Discovery Rate (FDR) correction was applied ^61^All tests are 2-sided. Data were analysed fully blind to condition.

**Response attrition at follow-up**

Attrition in response was seen at each in all groups at each follow-up time-point. *Table S1,* below gives the respondent Ns at each time point split by group.

Table S1: N respondents in each group at each time point from baseline to final follow up for all drinking-related measures.

|  |  | baseline | post-manipulation | 2 weeks | 3 months | 6 months | 9 months |
| --- | --- | --- | --- | --- | --- | --- | --- |
| AUDIT | No RET no PE | 22 | 30 | 27 | 25 | 25 | 26 |
|  | No RET+PE | 20 | 29 | 24 | 23 | 23 | 26 |
|  | RET no PE | 22 | 30 | 27 | 23 | 23 | 23 |
|  | RET+PE | 20 | 29 | 30 | 27 | 27 | 23 |
| TLFB | No RET no PE | 30 | 30 | 27 | 23 | 23 | 26 |
|  | No RET+PE | 30 | 30 | 24 | 21 | 22 | 25 |
|  | RET no PE | 30 | 30 | 27 | 23 | 23 | 23 |
|  | RET+PE | 30 | 30 | 29 | 26 | 26 | 23 |
| SOCRATES | No RET no PE | 30 | 30 | 27 | 25 | 26 | 26 |
|  | No RET+PE | 30 | 30 | 24 | 23 | 24 | 26 |
|  | RET no PE | 30 | 30 | 27 | 23 | 22 | 23 |
|  | RET+PE | 30 | 30 | 30 | 27 | 25 | 22 |
| ACQ | No RET no PE | 30 | 30 | 27 | 25 | 26 | 26 |
|  | No RET+PE | 30 | 30 | 24 | 23 | 24 | 26 |
|  | RET no PE | 30 | 30 | 27 | 23 | 22 | 23 |
|  | RET+PE | 30 | 30 | 30 | 27 | 25 | 22 |

**RESULTS:**

**Manipulation Checks:**

Variability in learning across the counterconditioning task as well as responses to the UCSs themselves was evident across the sample. Some participants showed very large reductions in *Beer Bit* and *Beer Pic CS* liking, while others showed *increases* in liking of these CSs across the task, despite clear pairing with aversive UCSs. Equally, while most participants rated the *Pic* and *Bitrex* as highly unpleasant, some rated the pictures as ‘extremely pleasant’ and some even rated the Bitrex above the median point on the scale (i.e. slightly pleasant). Central and dispersion statistics for these ratings are given in *Table S1*.

*Table S2:* Variability in responses to CSs and UCSs during counterconditioning. Response heterogeneity in ‘level’ of counterconditioning is evident in the range of liking ratings and standard deviation (SD).

|  | **Min** | **Max** | **Mean** | **SD** |
| --- | --- | --- | --- | --- |
| *Beer-Pic CS liking Trial 1* | 2.5 | 10 | 7.82 | 1.82 |
| *Beer-Pic CS liking Last Trial* | 0 | 10 | 6.51 | 3.07 |
| *Beer-Bit CS liking Trial 1* | 2.5 | 10 | 7.28 | 1.68 |
| *Beer-Bit CS liking Last Trial* | 0 | 10 | 5.83 | 2.71 |
| *Neut-Neut CS liking Trial 1* | 2.5 | 10 | 7.1 | 1.71 |
| *Neut-Neut CS liking last Trial* | 0 | 10 | 7.14 | 2.39 |
| *∆ Beer-Pic CS liking* | -9.5 | 4.5 | -1.3 | 2.67 |
| *∆ Beer-Bit CS liking* | -9 | 3 | -1.44 | 2.47 |
| *∆ Neut-Neut CS liking* | -9.5 | 5.5 | 0.05 | 2.25 |
| *Bitrex UCS liking Trial 1* | 0 | 6 | 1.58 | 1.57 |
| *Bitrex UCS liking Last Trial* | 0 | 6.5 | 1.25 | 1.63 |
| *Pic UCS liking Trial 1* | 0 | 10 | 1.58 | 1.85 |
| *Pic UCS liking Last Trial* | 0 | 10 | 1.61 | 1.89 |
| *Neut UCS liking Trial 1* | 0 | 10 | 5.23 | 2.04 |
| *Neut UCS liking Last Trial* | 0 | 10 | 5.76 | 2.15 |

*Table S3*: Pearson’s correlations between acute changes in liking of beer cues counter conditioned with Bitrex (Beer-Bit) and pictorial (Beer-Pic) UCSs with Day 3 cue and alcohol reactivity outcomes.

|  |  | **No RET No PE** | | **No RET + PE** | | **RET no PE** | | **RET+PE** | |
| --- | --- | --- | --- | --- | --- | --- | --- | --- | --- |
|  |  | ∆ Beer-BIT | ∆ Beer-PIC | ∆ Beer-BIT | ∆ Beer-PIC | ∆ Beer-BIT | ∆ Beer-PIC | ∆ Beer-BIT | ∆ Beer-PIC |
| **Cue image Ratings** | **Beer-React liking** | .089 | -.101 | -.188 | -.106 | -.113 | .21 | .178 | .212 |
|  | **Beer-Non-React liking** | -.086 | -.121 | -.1 | -.025 | -.084 | .161 | .185 | **.37*** |
|  | **Wine Liking** | .085 | .141 | -.023 | -.02 | -.075 | .278 | .275 | .322 |
|  | **OJ Liking** | .249 | -.131 | -.149 | -.20 | -.096 | -.117 | -.204 | -.071 |
|  | **Beer-React urge** | .043 | .073 | -.242 | -.161 | .249 | -.013 | .171 | .295 |
|  | **Beer-Non-React urge** | -.087 | -.142 | -.154 | -.099 | .192 | -.012 | .222 | .319 |
|  | **Wine Urge** | .08 | .184 | -.093 | -.026 | -.101 | -.315 | .134 | **.39*** |
|  | **OJ Urge** | .093 | -.155 | -.177 | -.083 | .225 | -.177 | .077 | **.38*** |
| **In vivo beer ratings** | **Drink itself liking** | -.016 | .066 | -.21 | -.173 | -.154 | .101 | .324 | .058 |
|  | **Drink itself urge** | .037 | .086 | -.314 | -.221 | -.262 | .155 | **.363*** | **.441*** |
|  | **anticipated enjoyment** | .137 | .074 | -.243 | -.188 | -.35 | .052 | .247 | **.436*** |
|  | **Anticipatory urge** | .046 | .046 | -.314 | -.159 | -.349 | -.006 | .31 | **.445*** |
|  | **Drink enjoyment** | .148 | .209 | -.026 | .027 | -.143 | -.073 | .305 | **.39*** |
|  | **Post-drink want more** | .161 | .098 | -.041 | .053 | -.016 | -.075 | .**367*** | **.515*** |

**Success of memory reactivation procedures**:

*Motivational impact of retrieval cues*:

*‘Liking’* of relevant drink cues (beer or orange juice) during the retrieval/no retrieval manipulation was assessed with *RET X PE X Cue Type* ANOVA. For this analysis, the liking ratings were averaged for the relevant ‘retrieval’ images (Beer images in RET groups and orange juice images in No RET groups) and for the ‘neutral’ drink cues (coffee and cola images in all groups). This revealed a main effect of *Cue Type* and a *Cue Type x Retrieval x PE* interaction [F(1,116) = 5.429, *p* =.024, *η_p_^2^*= .043]. Comparison of the simple effects of *Cue Type* within each group showed that the relevant reactivation cues were liked more than the neutral coffee/cola neutral cues in all groups (all F(1, 116) > 5.475, *p<*.021, *η_p_^2^*> .045) except for the *No RET + PE* group, where the orange juice images was not significantly greater than the cola/coffee images [F(1, 116) = 3.708, *p* = .057, *η_p_^2^* = .031]. No between-group differences were observed. ‘*Urge to drink*’ the relevant drink (beer in RET groups, or orange juice in No RET) in response to retrieval cues showed main effects of *Cue Type* [F(1, 116) = 123.075, *p<*.0001, *η_p_^2^* = .515] and *Retrieval* [F(1, 116) = 5.703, *p*=.019, *η_p_^2^* = .047]. In all groups, *urge to drink* was higher in response to the relevant retrieval cues than neutral drink (coffee/cola) cues. Cue-induced *urge to drink* beer in the *RET* groups was lower than cue-induced urge to drink orange juice in the *No RET* groups.

*Motivational impact of in-vivo drink reward*: Pre the prediction-error generation procedure, there were no group differences in liking of (*ps* >.719 *η_p_^2^s*<.001) anticipated enjoyment of (*ps* >.685 *η_p_^2^s*<.001) or urge to drink (*ps* >.719 *η_p_^2^s*<.001) the *in vivo* sample of beer or orange juice. In the *No PE* groups (where the drinks were actually consumed during retrieval) there was no group difference between actual enjoyment of the drinks [F(1,58) = .223, *p*=.639, *η_p_^2^* = .004] nor *desire to drink more* of the drink [F(1,58) = .142, *p*=.708, *η_p_^2^* = .003]. In total, this indicates that the *RET* and *No RET* procedures were well matched in terms of their ability to engage hedonic and motivational consumption processes.

*Prediction error generation:*  Withholding drink reward in PE groups is intended to induce cognitive prediction error or ‘*surprise*’. Analysis of rated ‘surprise’ levels following the retrieval and PE/no PE procedures showed a main effect of PE, indicating greater surprise following the PE procedure than the no PE procedure drink [F(1,116) = 309.79, *p*<.001, *η_p_^2^* = .728]. This did not interact with *Retrieval* group. The PE generation procedure was thus highly successful and equally effective in *RET* and *no RET* groups.

**Counterconditioning**

***Aversiveness of UCSs*:** A main effect of UCS Type (Bitrex > Disgusting Picture > neutral picture) was observed [F(2,230)=284.791, *p*<.0001, η_p_^2^= .712], along with a *UCS Type X Trial* interaction. The interaction indicated cumulative aversion in response to Bitrex UCS, with pleasantness ratings becoming more extremely negative across *Trials* [Trial simple effect for Bitrex F(3,113) = 5.712, *p* = .001, η_p_^2^= .132]. There were no effects or interaction with *Retrieval* or *PE* groups. *Overall*, the disgusting UCSs were thus effective negative reinforcers during counterconditioning.

**Predictors of response to counterconditioning and changes in drinking.**

Disgust propensity and sensitivity were predictive of alcohol consumption during the post-manipulation period, with greater general propensity to disgust [*r*(120) = -.31, *p* = .001] and sensitivity to disgusting stimuli [*r*(120) = -.365, *p<*.001] predicting lower total alcohol consumption. Disgust propensity was also associated with participants’ mean ratings of the unpleasantness of the disgusting images during counterconditioning, indicating higher rated unpleasantness with greater disgust propensity [*r*(120) = -.311, *p* = .001]

Pleasantness ratings of the Bitrex UCSs were negatively associated with post-manipulation AUDIT scores [*r*(117) = -.259 *p* = .005] and urge to drink in response to beer images [*r*(119) = -.213 *p* = .005]. In-lab ratings of reactivity were moderately, (but significantly) correlated with questionnaire-measured craving and drinking outside of the lab (*rs 0.2 – 0.39, p*s 0.01-0.029).

Peri-reactivation affect and arousal may be key moderators of counterconditioning effects, since counterconditioning is an inherently aversive procedure which may interact with negative affect and anxiety in strengthening learning. Further, emotional arousal is well established to potentiate associative learning. Indeed, *arousal* induced by exposure to drug stimuli without reinforcement has been posited as a possible explanation for the enhancing effect of retrieval-extinction procedures, rather than memory rewriting ^36^.

In support of this interpretation, pre-counterconditioning state anxiety levels on the STAI modestly negatively predicted beer total drinking levels [*r*(120)=-.268, *p*=.003] post-manipulation and at 2-week follow up [*r* (107) = -.214, *p* =.027], but not 3 months, 6 months or 9 months. Similarly, negative affect, derived from the PANAS predicted lower drinking levels at post-manipulation [*r* (120) = -.188, *p* =.04] and 2 weeks [*r* (107) = -.212, *p* =.029] but not longer-term follow ups periods. This is consistent with the engagement of dual processes; affective potentiation of counterconditioning (new learning), yielding shorter-term effects on maladaptive drinking behaviour, with a reconsolidation-based *rewriting* mechanism accounting for more durable long-term reductions in drinking.

***Exploratory subgroup analysis of ‘responders****’*: Analysis of only participants who were responsive to counterconditioning (defined as those who reduced their liking of *Beer-Pic* AND *Beer-Bit* cues from the first to last trial of counterconditioning, as is common in conditioning literature), yielded group *N*s of No *RET no PE* =15, *No RET+PE*=10, *RET no PE*=10, *RET+PE*=15. Thus only half, or fewer, of participants acutely displayed ‘full’ counterconditioning of cues. Re-analysis of reactivity to the beer with *Day X RET X PE* ANOVAs in these groups revealed trend-level *Day*RET*PE* interactions for *urge to drink* [F(1,46)=3.17, *p*=.082, *η_p_^2^*= .064]. Multivariate simple effects analyses revealed that this was due to an effect of *Retrieval* in the *PE groups* on *Day 3*, representing lower *urge to drink* in *RET+PE* than *No RET+PE* [F(1,46)=5.281, *p*=.026, *η_p_^2^*= .103].

**Cue reactivity: Responses to cue images**

Ratings of cue image pleasantness and urge to drink depicted beverages during the cue reactivity task were assessed by *Day* (Baseline, post-manipulation) x Retrieval (RET vs No RET) x Prediction Error (PE+/PE-) x Cue Type (Reactivated beer, Non-reactivated beer, wine, orange juice, soft drink) mixed ANOVA, with counterconditioning responsiveness included modelled as a covariate in a fully factorial model.

**Urge to drink depicted beverages** A *Cue Type* main effect [multivariate F(4,112)=49.353, *p*<.001, *η_p_^2^*=.638] and *Day*Cue Type* interaction [multivariate F(4,112)=7.059, *p<*0.001, *η_p_^2^*=.201] were found, subsumed under a *Retrieval*PE*Day*Cue Type* interaction [multivariate F(4,113)=3.823, *p* =.006, *η_p_^2^*= .12]. The latter interaction was investigated by splitting the analysis by *RET* vs *No RET* groups. A *Day*Cue Type*PE* interaction was present only in the *RET* groups [F(2.424, 191.915)=3.615, *p*=.011, *η_p_^2^*= .06]. Inspection of the simple multivariate effects of *Day* indicated that *RET+PE* displayed a significant *reduction* in induced *urge to drink* depicted beer in response to reactivated/ counterconditioned beer cues (*Beer RET*; F(1,57)= 5.5856, *p*=.019, *η_p_^2^*=.093) and a significant *increase* in urge to drink orange juice [F(1,57)= 7.293, *p* =.009, *η_p_^2^*= .113]. Conversely, in *Ret no PE*, decreases were seen in urge to drink reactivated beer cues [F(1,57)=4.659, *p*=.035, *η_p_^2^*=.076] and wine cues F(1,57)=5.771, *p*= .02, *η_p_^2^*=.092].

*Figure S1*: Effects of counterconditioning in *RET* groups on *urge to drink* depicted beverages post-manipulation in the cue reactivity task. BR = reactivated beer images, BNR = Non-reactivated beer images, Wine = Wine images, OJ = Orange juice images, Neutral=coffee/cola images. Bars represent mean±SEM

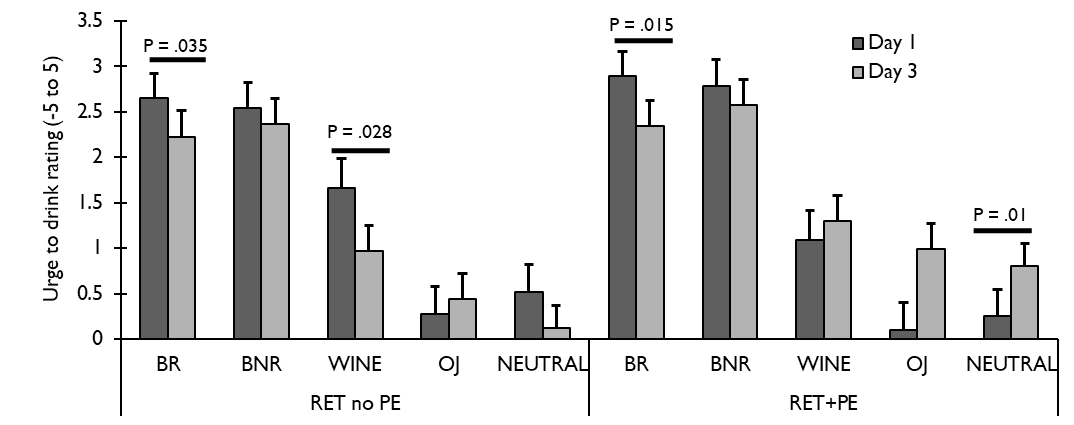

**Liking of cues:**

A main effect of *Cue Type* [*p<*0.001 *η_p_^2^*=.164) was found, subsumed under a *Day*Cue Type*Responsiveness* interaction [F(3.351,385.39)=3.905, *p*=.007, *η_p_^2^*=.033]. Analyses on each *Cue Type* showed *Day*Responsiveness* interactions for reactivated beer images [F(1, 115)=4.673, *p* =.033, *η_p_^2^*=.039] and non-reactivated beer images [F(1, 115)=4.665, *p*=.033, *η_p_^2^*=.039], but not wine, orange juice or soft drink (neutral) images. For both types of beer image, greater counterconditioning responsiveness predicted lower *Day 3* liking.

**Follow-up data secondary measures**

**Craving (ACQ-NOW)**

General self-rated craving according to the ACQ-NOW did not change in the short-term between pre-and post-manipulation [F(1,116)=1.19, *p*=.278, η_p_^2^=.01], however mixed-model analysis with random slopes for *Time* showed long-term reduction in craving over the follow-up period up to 9 months [F(1,106.194)=260.895, *p<*0.001]. Contrasts on estimated marginal means demonstrated significant reductions in craving by the 2 week follow up [F(1,82)=87.98, *p* <.001, *η_p_^2^*=.518] that persisted or further reduced at all follow-up time points up to at least 9 months [F(1,82)=284.9, *p* <.001, η_p_^2^=.777].

**Readiness to Change (SOCRATES):** All groups reported greater recognition of the need to change their drinking behaviour [F(1, 116)=7.378, *p*=.008, η_p_^2^=.06], reductions in ambivalence towards their excessive drinking [F(1, 116)=8.897, *p*=.003, η_p_^2^=.071] and increases in ‘taking steps’ to reduce their drinking [F(1, 116)=16.11, *p*<.001, η_p_^2^=.122], from baseline to post-manipulation. These beneficial changes did not differ according to *RET* or *PE* group.
